# Laplacian eigenmaps and principal curves for high resolution pseudotemporal ordering of single-cell RNA-seq profiles

**DOI:** 10.1101/027219

**Authors:** Kieran Campbell, Chris P Ponting, Caleb Webber

## Abstract

Advances in RNA-seq technologies provide unprecedented insight into the variability and heterogeneity of gene expression at the single-cell level. However, such data offers only a snapshot of the transcriptome, whereas it is often the progression of cells through dynamic biological processes that is of interest. As a result, one outstanding challenge is to infer such progressions by ordering gene expression from single cell data alone, known as the cell ordering problem. Here, we introduce a new method that constructs a low-dimensional non-linear embedding of the data using laplacian eigenmaps before assigning each cell a pseudotime using principal curves. We characterise why on a theoretical level our method is more robust to the high levels of noise typical of single-cell RNA-seq data before demonstrating its utility on two existing datasets of differentiating cells.

## 1 Introduction

Single-cell RNA-seq (scRNA-seq) is a powerful method for the quantification of transcript abundance in individual cells. It has already led to new discoveries in biology including the identification of novel cell types [1], hidden heterogeneity in gene expression [2] and cellular regulatory networks [3]. It has distinct advantages over bulk RNA sequencing where expression estimates are averaged over all cells, masking features of interest such as cell type specificities, intrapopulation heterogeneity and gene (co)expression [4] [5]. However, as transcriptional networks are dynamic each individual cell’s transcriptome will be reflecting its progression through many biological processes, such as differentiation or apoptosis [6]. Consequently, we would like to assign to each cell an estimate of its progression (known as its *pseudotime*) in a hypothesis-free data-driven manner (Supplementary Figure 1). Subsequently, gene expression dynamics across pseudotime, such as differential expression and clusters of co-expression, can be inferred.

The current leading method in the field for RNA-seq data is *Monocle* [6]), which uses independent component analysis (ICA) to reduce the dimension of the problem before fitting the pseudotime in the reduced space using the longest path through a minimum spanning tree (MST). However, ICA can only reduce the dimensionality to linear combinations of genes while the fastICA algorithm [7] it uses is highly sensitive to outliers [8] that are typical of scRNA-seq datasets [9]. Furthermore, fitting pseudotime using a graph-based as opposed to curve-based approach may fit to some of the noise inherent in the dataset. Further methods used for pseudotime ordering include *Wanderlust* [10] for mass-cytometry data, *Waterfall* which connects connects k-means clusters in PCA space [11] and an approach that uses diffusion maps for visualisation and imputation [12].

Here we present *Embeddr* (available at http://www.github.com/kieranrcampbell/embeddr as an R package), a method for the pseudotemporal ordering of scRNA-seq data that is robust to the typically high levels of technical and biological noise and requires no prior knowledge of phenotypic marker genes. *Embeddr* employs non-linear dimensionality reduction of the gene-space using *laplacian eigenmaps* (also known as *spectral embedding*) [13] [14] - a manifold learning technique - before tracing through the centre of the manifold using principal curves [15] and assigning pseudotime based on arc-length from the edge of the manifold. We apply *Embeddr* to two existing scRNA-seq datasets and show that while it is consistent with current methods, it is more sensitive to cell type identification and may be used to identify marker genes across pseudotime.

## 2 Methods

In scRNA-seq data cells are represented by gene-expression vectors (20,000+ dimensions depending on whether individual transcripts are considered) that lie in a high-dimensional space. However, if the main source of variation in the dataset is a biological process of interest that is captured in the cell’s transcriptomic profiles then the cells will not be randomly scattered through this transcriptomic space but instead will lie on a lower-dimensional noisy surface known as a manifold. Thus, traversing through this manifold would track the cells as their gene expression evolves through pseudotime.

To reveal this manifold we use laplacian eigenmaps that, given a cell-to-cell similarity matrix *W*_*ij*_, attempt to minimise the quantity

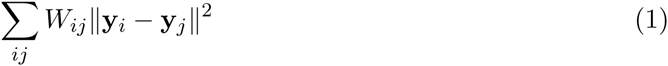

where **y**_*i*_ is the *p*-dimensional embedding of cell *i*. In other words, if two cells are transcriptionally similar we attempt to place them nearby in the low-dimensional embedding. It can be shown that minimisation of equation 1 subject to the constraint **y**^*T*^ **y**= 1 is equivalent to solving the eigenvalue equation

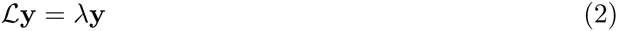

where *ℒ* = *D − W* and *D* is the *degree matrix* and is the diagonal matrix made up of the row sums of *W*. The multiplicity of the 0 eigenvalue of the laplacian matrix *ℒ* corresponds to the number of connected components in the graph *W*. If there are more than one connected component in the graph then the embedding is highly unstable and should be re-fitted for each component. However, there is great utility to seeing the relationship between different cells on a single plot, so the user may wish to increase the number of nearest neighbours *n* until a single connected component is obtained.

We can created a non-linear mapping that is robust to noise through the choice of *W*. First, a cell-by-cell correlation matrix across high-variance genes is constructed. Then, using this as a measure of similarity, *W* is defined as

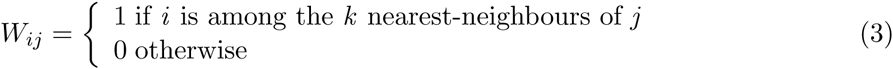

where *k* is typically chosen as *k* = ln(*n*) for *n* cells. When *n* is relatively small (*<*100) it can be advantageous to select *k >* log(*n*) because if *k* is very small the resulting embedding may be unstable. The specification of *W* encourages a non-linear embedding as traversing the manifold is equivalent to passing through successive nearest neighbours, allowing the global structure to be warped. Furthermore, because cell transcriptomic similarity is only defined as being one of *k* nearest neighbours or not, laplacian eigenmaps can learn noisy manifolds [13] as outliers typical of the data will have minimal effect. Somewhat counterinuitively, cell *i* being among the *k* nearest neighbours of cell *j* does not imply cell *j* is among the *k* nearest neighbours of *i*. Since the minimisation in equation 1 is achieved by solving an eigenvalue equation in *W* we instead use the symmetrised weight matrix 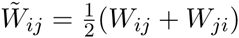to ensure that the eigenvalues are real and the eigenvectors orthogonal.

Once this low (typically 2) dimensional embedding is constructed, principal curves [15] are used to trace through the centre of the manifold and assign a pseudotime to each cell. Principal curves are smooth, non-parametric curves that pass through the middle of a dataset and whose shape is entirely determined by the data. They are preferable to standard curve-fitting algorithms such as least squares regression because they require no model choice (e.g. the degree of the polynomial), they may be multi-valued and are agnostic under axis switching.

Once the principal curve has been fit, the orthogonal projection of each cell onto it may be found (Supplementary Figure 2). Subsequently, the arc-length from the beginning of the curve to the projected point estimates the pseudotime of the cell. All pseudotimes are then rescaled to be in [0,1] by 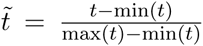. By projecting each cell into the centre of the manifold an additional source of orthogonal noise is minimised which may be present in graph-based pseudotime approaches.

The overall workflow for *Embeddr* is shown in figure 1. First, high-variance genes are selected using spike-ins if available or by fitting coefficient of variation - mean curves in a manner similar to [16]. The entire gene-set may be used providing similar results (Supplementary Figure 3) or only a given subset of genes, for example those genes within a pathway of interest whose variation might be masked by other high-variance genes. Once the laplacian eigenmaps embedding has been found it can be clustered using unsupervised algorithms such as k-means or Gaussian mixture models to remove outlying cells for a given pseudotime trajectory, such as contaminating cells or those on a separate trajectory. The pseudotime ordering is then found as above and differential expression and co-expression analysis can be performed in a similar manner to previous work [6]. *Embeddr* combines aspects of both Monocle and *Wanderlust* by first performing dimensionality reduction as with Monocle, which is helpful for visualisation, but using an underlying *k*-nearest-neighbour graph as the similarity matrix like Wanderlust, which allows for non-linear embeddings.

**Figure 1:**
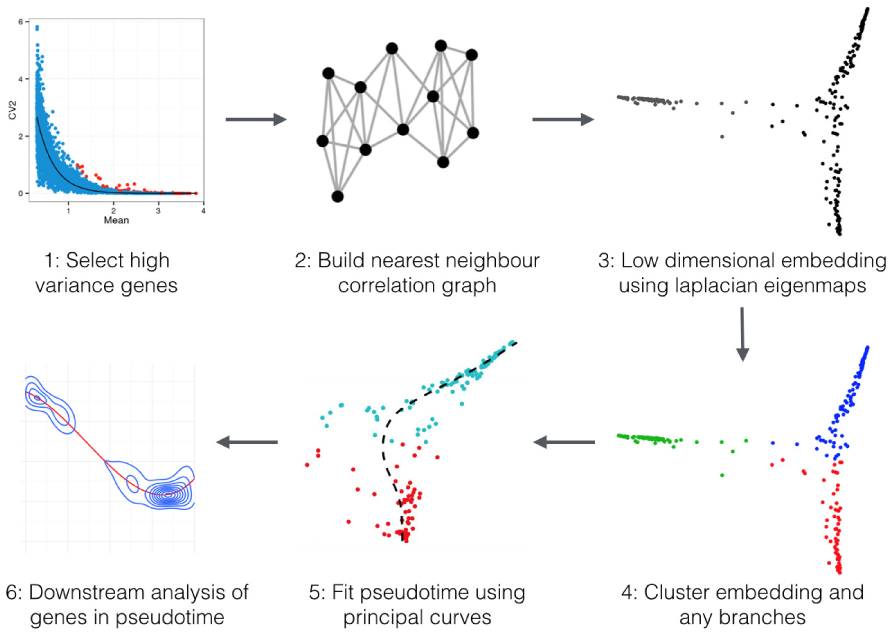
*Embeddr* workflow. (1) Optionally select high-variance genes using spike-ins (if available) or by fitting coefficient of variation - mean curves. (2) Build nearest neighbour graph based on correlation metric between cells, typically choosing *k* = log(*n*) nearest neighbours for n cells. (3) Find the low-dimensional laplacian eigenmaps embedding of the cells using the nearest-neighbour graph. (4) Cluster the embedding using k-means, Gaussian mixture models or any other appropriate unsupervised clustering algorithm. (5) Fit the pseudotime along a given trajectory using principal curves. (6) Analysis of genes through pseudotime including differential expression and clusters of co-expression.

## 3 Results

We applied our method to two existing single-cell RNA-seq datasets: a set derived from differentiating human myoblasts [6] and a further set derived from differentiating cells in the mouse distal lung epithelium [1].

### 3.1 Application to differentiating human myoblasts

Our first application of *Embeddr* was to scRNA-seq data derived from primary human skeletal muscle myoblasts that had been expanded in high-mitogen conditions before being differentiated in a low-serum medium. Subsequently, transcriptomes for 271 single cells were sequenced at 4 time points. The original analyses inferred highly asynchronous differentiation with myoblasts, intermediate myocytes and mature myotubes present at each time point, thereby exemplifying the need for a pseudotemporal ordering of cells.

The Laplacian Eigenmap embedding of the cells was found using *Embeddr* (figure 2), showing a clear differentiation trajectory amongst one group of cells and identifying a separate cluster of contaminating cells. We next plotted several marker genes as employed in the original paper to discern different cell types (figure 2), whose expression clearly separates those cells undergoing differentiation from the (contaminating) mesenchymal cells. We compared our classification of cell types to that of *Moncole* and found we classified substantially more cells as mesenchymal (116 vs 61, Supplementary Table 1). We found high expression of mesenchymal marker genes in cells designated as mesenchymal by *Embeddr* but as proliferating/differentiating by Moncole (Supplementary Figure 4), indicating that the application of laplacian eigenmaps provides increased sensitivity of cell type classification.

**Figure 2:**
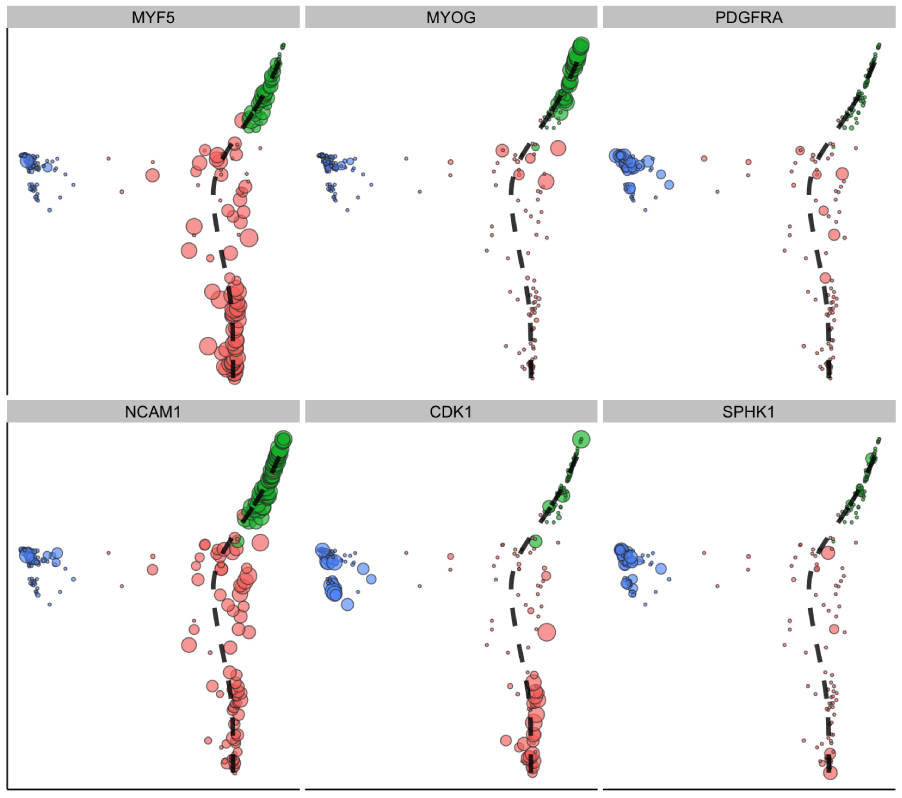
Expression of marker genes in laplacian eigenmaps embedding with the black dashed pseudotime fit. *MYF5* and *NCAM1* (left) are markers of proliferating/differentiating cells, suggesting the set of cells on the right are those undergoing differentiation. *CDK1* (bottom, middle) is a marker gene for early proliferating cells while *MYOG* (top, middle) is a marker for late differentiating cells, suggesting the differentiation trajectory runs from bottom (red) to top (green). Finally, *PDGFRA* and *SPHK1* (right) are markers for the (contaminating) mesenchymal cells, providing further support to the notion that the blue cluster are the mesenchymal cells while the red and green are early and late differentiation respectively.

Considering only those cells classified as proliferating or differentiating we fitted the pseudotime of each cell using principal curves, as described above (Supplementary Figure 5). This displayed high correlation with the pseudotime assigned using *Moncole* (Spearman correlation coefficient 0.8, Supplementary Figure 6). We then identified genes differentially expressed across pseudotime using tobit regression and smoothing splines as previously described in [17], and clustered them using complete linkage clustering with a distance metric *d*(*x, y*) = 1 *− ρ*_*xy*_/2. Hierarchical clustering identified 4 major clusters (Supplementary Figure 7) and enriched GO terms were identified for each cluster using Goseq [18]. Two of these clusters identified early down-regulation of cell-cycle and gradual up-regulation of muscle structure development respectively (Supplementary Table 2), which is consistent with myoblast differentiation.

We validated the robustness of the pseudotemporal ordering to both parameter choice and selection of cells. First, we varied the number of nearest neighbours used in the construction of the correlation graph W and refitted both the embedding and the pseudotime (Supplementary Figure 8). In each case the clusters remained well separated and the principal curve fitted consistently. Next, we randomly removed 50% of cells 30 times and refitted the pseudotime (Supplementary Figure 9). As before, original clusters remained well separated and the pseudotimes showed excellent agreements between refits (Supplementary Figure 10, median Spearman correlation 0.98, first and third quartiles 0.97, 0.98 respectively). Finally, we sub-sampled 50% of the genes used for the embedding and refit the pseudotime, an approach previously suggested to provide statistical support to an ordering [19]. This showed a robust ordering of cells with high correlations between subsamples (Supplementary Figure 11, median Spearman correlation 0.91, first and third quartiles 0.86, 0.94 respectively).

### 3.2 Application to differentiating cells in the distal lung epithelium

We then applied our method to a second scRNA-seq dataset consisting of 80 cells profiled from the developing distal lung epithelium in mouse embryos. This dataset contains five cell types as identified by the authors, including bipolar progenitor (BP) cells, which differentiate into either alveolar type 1 (AT1) or alveolar type 2 (AT2) cells. The laplacian eigenmaps representation implied a continuous developmental trajectory between AT1, BP and AT2 cells for which the principal curve was used to fit pseudotime (figure 3). We then plotted the density of cells through pseudotime (Supplementary Figure 12) which clearly shows BP cells reside in a low heterogenous density state before moving into high density more homogeneous differentiated states. This implies that the progenitor cells have greater heterogeneity in their expression profiles before specialising into differentiated cells that are highly transcriptionally homogeneous.

**Figure 3:**
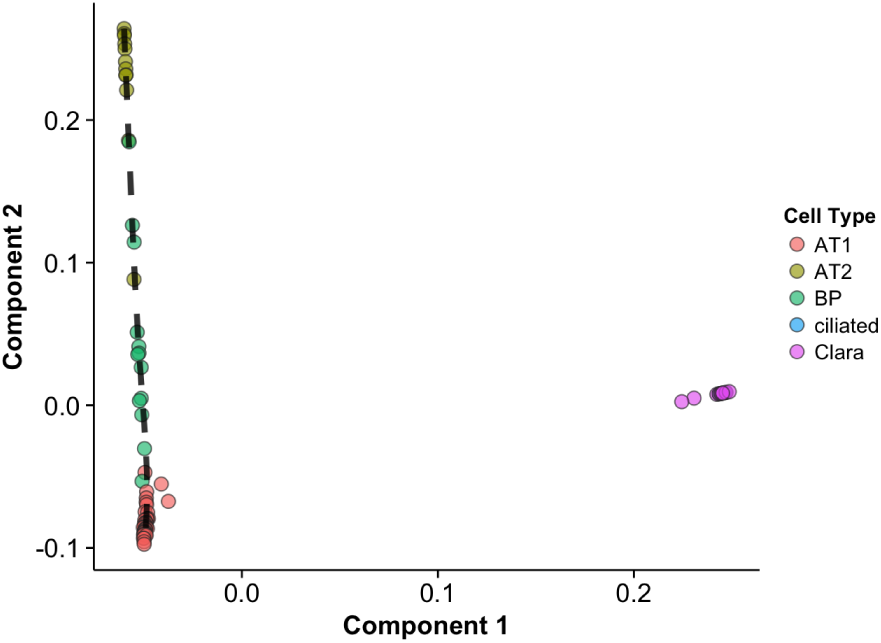
Laplacian eigenmaps embedding of distal lung epithelium data from Quake et al. Cell types are coloured by classification in the original paper, with the black dashed like showing the pseudotime fit through the differentiation trajectory. Cells of type BP differentiate into either AT1 or AT2 by selective silencing of genes. More mature Clara and Ciliated cells appear separate on the right hand side. *k* = 10 nearest neighbours were used in the similarity graph as a low number of starting cells were present.

Next we posited that differential expression of genes over pseudotime could be used to recover marker genes for the cell types present. The original analyses by [1] identified ‘perfect marker genes’ as those expressed at greater than 2^10^ FPKM in all cells of a given type and at 0 FPKM in all other cells. While this on-off method will identify easily discoverable transcripts in assays, it requires setting a hard threshold and ignores more gradual gene expression changes that are typical of differentiation. We refit our pseudotime trajectories for the BP to AT1 and BP to AT2 trajectories separately and performed differential expression tests for genes silenced across pseudotime using those that were expressed in more than 20% of BP-AT1 cells above 0 FPKM. A total of 28 genes were found to be differentially expressed across the BP-AT1 transition and 111 across the BP-AT2 transition. By examining clusters of cells switching on and off we recovered 11 of the 14 marker genes for AT1 cells and 15 of the 19 for AT2, and further identified 28 putative marker genes for AT1 and 76 for AT2 (Supplementary Figure 15, Supplementary Tables 3 and 4).

## 4 Discussion

We have presented a novel method for pseudotemporal ordering of single-cell RNA-seq profiles. By reducing the dimensionality of the dataset using laplacian eigenmaps we can discover highly non-linear embeddings of the observed transcriptomic space that are robust to high levels of noise. The progression of cells through pseudotime may then be defined by fitting a principal curve through the manifold identified. This allows us to identify gene dynamics such as differential expression and clusters of co-expression across pseudotime as well as novel marker genes for cell types involved in pseudotemporal processes.

Our method discovers a pseudotemporal ordering that is similar to previous methods (Supplementary Figure 4) but which cleanly separates out different cell types as identified by marker genes (figure 5) implying a pseudotemporal ordering with higher sensitivity. Our method combines many of the strengths of current methods such as the graph based Wanderlust and ICA-based Monocle, being able both to find non-linear embeddings defined by a flexible set of similarity metrics (such as correlation, cosine, Euclidean) and enable visualisation of cell types through operating in a reduced-dimension space.

How can pseudotime be interpreted and what is being discovered using *Embeddr* and similar methods? While in obvious cases such as the differentiating myoblasts pseudotime clearly corresponds to differentiation status, it may in other datasets simply correspond to any source of biological (or in unfortunate cases technical) variation present. In theory one could choose the set of genes corresponding to a particular biological pathway of interest and the resulting pseudotime ordering of cells will correspond to the extent of pathway progression. It should be noted that *Embeddr* will always be able to find a pseudotime ordering whether or not there is a real biological process to be modelled. Since, for sufficiently large n, a cell will always have a nearest neighbour, unless there are highly distinct cell types (which almost never happens since there are usually underlying pathways that induce correlations) then the low dimensional embedding can always be found and a pseudotime fitted. While this is useful if the user *a priori* suspects that there is an underlying pseudotemporal process, the fact that one may be fitted using the algorithm does not imply that such a process exists.

## Acknowledgement

The authors would like to thank Christopher Yau (University of Oxford) for his feedback on the manuscript.

